# Scanpy for analysis of large-scale single-cell gene expression data

**DOI:** 10.1101/174029

**Authors:** F. Alexander Wolf, Philipp Angerer, Fabian J. Theis

## Abstract

We present Scanpy, a scalable toolkit for analyzing single-cell gene expression data. It includes preprocessing, visualization, clustering, pseudotime and trajectory inference, differential expression testing and simulation of gene regulatory networks. The Python-based implementation efficiently deals with datasets of more than one million cells and enables easy interfacing of advanced machine learning packages. Code is available from https://github.com/theislab/scanpy.

---

Simple integrated analysis workflows for single-cell transcriptomic data (Stegle *et al.*, 2015) have been enabled by frameworks such as Seurat (Satija *et al.*, 2015), MAST (Finak *et al.*, 2015), Monocle (Trapnell *et al.*, 2012), Scater (McCarthy *et al.*, 2017), Cell Ranger (Zheng *et al.*, 2017), Scran (Lun *et al.*, 2016) and SCDE (Kharchenko *et al.*, 2014). However, they do not scale to the increasingly available large-scale datasets with up to one million cells. Here, we present a framework that overcomes this limitation and provides similar analysis possibilities (Fig. 1a). In addition, in contrast to the existing R-based frameworks, Scanpy’s Python-based implementation allows to easily integrate advanced machine learning packages, such as Tensorflow (Abadi *et al.*, 2015, Suppl. Note 1).

**Figure 1 | a.**
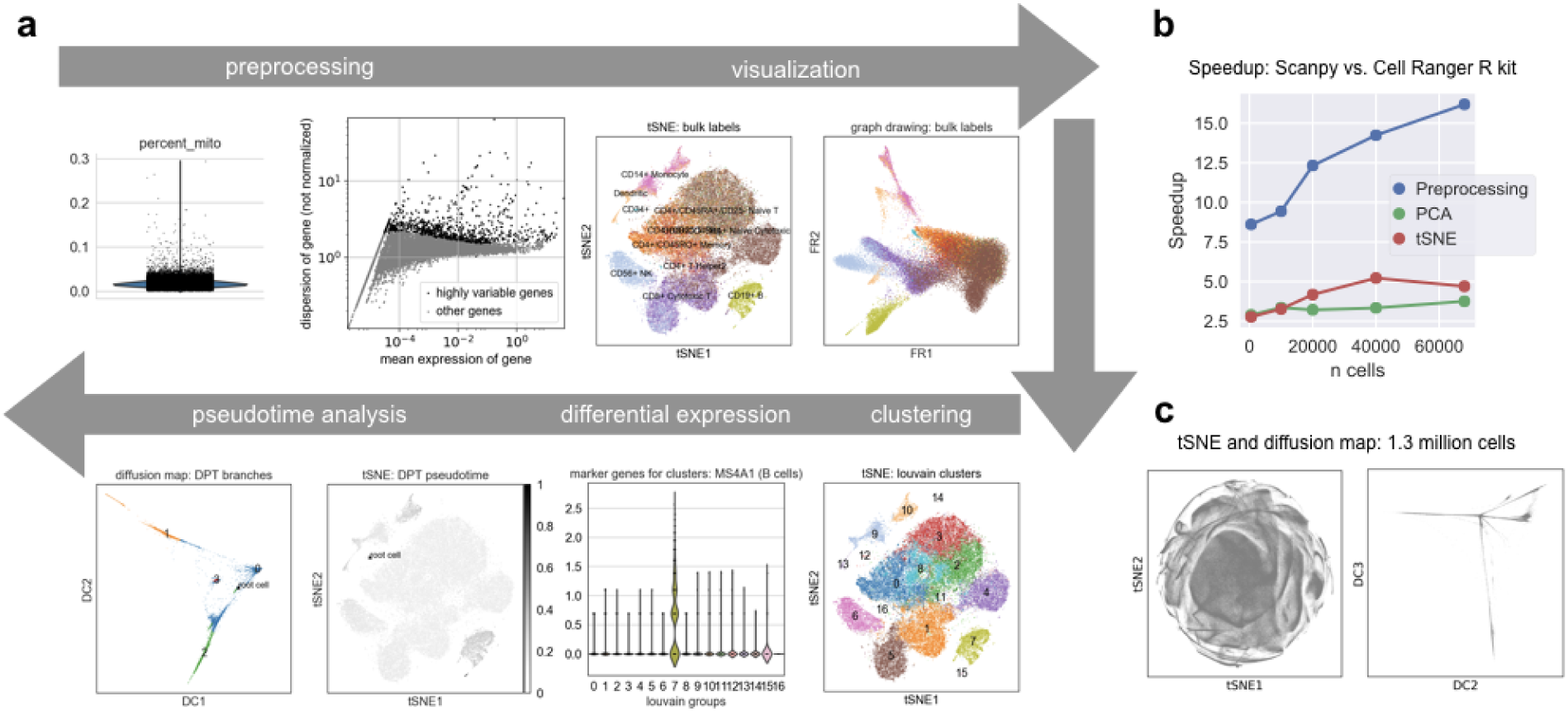
Scanpy’s analysis features. Using the example of 68,579 PBMC cells of Zheng *et al.* (2017). Regressing-out confounding variables, normalizing and identifying highly-variable genes. tSNE and graph-drawing (Fruchterman-Reingold) visualizations show cell types obtained by comparing with bulk expression (Zheng *et al.*, 2017). Louvain clustering. Ranking differentially-expressed genes in clusters identifies the MS4A1 marker gene for B cells in cluster 7, which agrees with the bulk labels. Pseudotemporal ordering from a root cell in the CD34+ cluster and detection of a branching trajectory, visualized with tSNE and Diffusion Maps. **b, Speedup over Cell Ranger R kit**. For representative steps of the analysis (Zheng *et al.*, 2017). **c, Visualizing 1.3 million cells**. Using tSNE and Diffusion Maps. The data, brain cells from E18 mice, is publicly available from 10X Genomics.

Scanpy provides preprocessing comparable to Seurat (Macosko *et al.*, 2015) and Cell Ranger (Zheng *et al.*, 2017) and visualization through tSNE (Coifman *et al.*, 2005; Amir *et al.*, 2013), graph-drawing (Fruchterman and Reingold, 1991; Csardi and Nepusz, 2006; Weinreb *et al.*, 2017), Diffusion Maps (Coifman *et al.*, 2005; Haghverdi *et al.*, 2015; Angerer *et al.*, 2015) and principal component analysis (Fig. 1a). It provides clustering similar to Phenograph (Blondel *et al.*, 2008; Levine *et al.*, 2015) and allows identifying clusters with cell types by finding marker genes using differential expression testing. Scanpy provides pseudotemporal-ordering and the reconstruction of branching trajectories via Diffusion Pseudotime (DPT, Haghverdi *et al.*, 2016)^1^ and allows simulating single cells governed by gene regulatory networks (Suppl. Note 2, Wittmann *et al.*, 2009). Scanpy provides its tools with speedups between 4 and 16 and much higher memory efficiency (about a factor 10) than comparable frameworks (Fig. 1b, Suppl. Note 2). This enables the analysis of datasets with over a million cells and allows an *interactive* analysis of about hundred thousand cells (Fig. 1c, Suppl. Note 2).

Scanpy is implemented in a highly modular fashion and can hence be easily further developed by a community (Suppl. Note 3). Its data storage formats and objects allow a simple cross-language and cross-platform transfer of results (Suppl. Note 3). Scanpy integrates well into the existing Python ecosystem, where no comparable toolkit yet exists (Suppl. Note 4).

## Acknowledgements

We thank the authors of Seurat for sharing their great tutorials. We thank S. Tritschler, L. Simon, D. S. Fischer, F. Buettner and G. Eraslan for commenting on the software package and the paper. F.A.W. acknowledges support by the Helmholtz Postdoc Programme, Initiative and Networking Fund of the Helmholtz Association.

## Supplemental Note 1: Scanpy’s technological foundations

Scanpy is on the Python packaging index: https://pypi.python.org/pypi/scanpy.

Scanpy’s Python-based implementation allows easily interfacing advanced machine learning packages such as Tensorflow (Abadi *et al.*, 2015) for Deep Learning (LeCun *et al.*, 2015), Limix for linear mixed models (Lippert *et al.*, 2014) and GPy/GPflow for Gaussian Processes (GPy, 2012; Matthews *et al.*, 2017). See Suppl. Note 2 for an example of combining Deep Learning and Scanpy (Eulenberg *et al.*, 2016).

Scanpy’s core relies on Numpy (van der Walt *et al.*, 2011), Scipy (Jones *et al.*, 2001), Matplotlib (Hunter, 2007) and h5py (Collette, 2013). Parts of the toolkit rely on Pandas (McKinney, 2010), scikit-learn (Pedregosa *et al.*, 2011), statsmodels (Seabold and Perktold, 2010), Seaborn (Waskom *et al.*, 2016), NetworkX (Hagberg *et al.*, 2008), igraph (Csardi and Nepusz, 2006), the tSNE package of Ulyanov (2016) and the Louvain clustering package of Traag (2017).

## Supplemental Note 2: Scanpy’s analysis features

The following links allow to reproduce Figure 1, give detailed background information on the benchmark computations and provide further use cases.

- The analysis of 68,579 PBMC cells of Figure 1 and the comparison with the Cell Ranger R kit (Zheng *et al.*, 2017): [scanpy_usage/170503_zheng17](https://github.com/theislab/scanpy_usage/tree/master/170503_zheng17).
- A detailed clustering tutorial, adapted from Seurat’s tutorial, walks the user from raw data through all steps of the analysis to the identification of cell types: [scanpy_usage/170505_seurat](https://github.com/theislab/scanpy_usage/tree/master/170505_seurat).
- Visualizing 1.3 mio cells as in Figure 1c: [scanpy_usage/170522_visualizing_one_million_cells](https://github.com/theislab/scanpy_usage/tree/master/170522_visualizing_one_million_cells).
- Examples for reconstructing branching processes via Diffusion Pseudotime (Haghverdi *et al.*, 2016): [scanpy_usage/170502_haghverdi16](https://github.com/theislab/scanpy_usage/tree/master/170502_haghverdi16).
- Simulating single cells using gene regulatory networks (Wittmann *et al.*, 2009); here, myeloid differentiation (Krumsiek *et al.*, 2011): [scanpy_usage/170430_krumsiek11](https://github.com/theislab/scanpy_usage/tree/master/170430_krumsiek11).
- Analyzing deep learning results for single-cell images (Eulenberg *et al.*, 2016): [scanpy_usage/170529_images](https://github.com/theislab/scanpy_usage/tree/master/170529_images).

## Supplemental Note 3: Scanpy’s technological concepts

Scanpy tools operate on a class *AnnData*, which simply stores the annotated data matrix. While Scanpy is in large parts object oriented, by building *AnnData*, we chose a functional-programming oriented design to enable a modular development of Scanpy: adding new functionality to the toolkit is easy as any new tool leaves the structure of *AnnData* unaffected. *AnnData* is similar to R’s ExpressionSet (Huber *et al.*, 2015), but supports sparse data and file iterators and provides simple control of the underlying data types (van der Walt *et al.*, 2011). In addition, *AnnData*’s simple structure allowed us to design a corresponding *hdf5* file format (Collette, 2013), which enables writing and reading objects to disk in a highly efficient and platform-, framework- and language-independent way. This allows easily transferring data and analysis results from and to existing R packages (see also Suppl. Note 3).

Further technological concepts are as follows.

- Support of reading a wide variety of data file formats and their simple cache in fast *hdf5* files; similar to caching full *AnnData* objects.
- A central class *DataGraph* whose focus is the efficient representation of a graph of neighborhood relations in data; their computation is parallelized and much faster than in existing packages (Pedregosa *et al.*, 2011). The class provides functions to compute quantities on the graph, which are not available in other graph packages (Hagberg *et al.*, 2008; Csardi and Nepusz, 2006). Storage is again platform- and language-independent via CSR sparse matrices, which appear as data annotation in *AnnData*.
- Scanpy functions by default operate “inplace” and thereby encourage and enable easily building memory-efficient pipelines.
- Computations are monitored by profiling information so that users develop an intuition for waiting times. In addition, this encourages performance-aware development.
- A modular design of the toolkit with user submodules for *preprocessing*, *tools*, *plotting*, *settings* and correspondence in naming conventions between the modules.
- A command-line interface that parallels the usage of the API allows easily submitting jobs to remote computing infrastructure.

Just before submission of this manuscript we became aware of an alternative approach to tackling large-scale data in statistical computing. Lun *et al.* (2017) provide a C++ library that simplifies interfacing large-scale matrices for R-package developers. This approach is therefore an alternative to only a small subset of Scanpy’s features — interfacing hdf5-backed large-scale matrices.

## Supplemental Note 4: Python packages for single-cell analysis

Aside from the highly popular [scLVM](https://github.com/PMBio/scLVM) (Buettner *et al.*, 2015, 2016), which uses Gaussian Process latent variable models for inferring hidden sources of variation, there are, among others, the visualization frameworks [FastProject](https://github.com/YosefLab/FastProject) (DeTomaso and Yosef, 2016), [ACCENSE](http://www.cellaccense.com/) (Shekhar *et al.*, 2013) and [SPRING](https://github.com/AllonKleinLab/SPRING) (Weinreb *et al.*, 2017),^2^ the trajectory inference tool [SCIMITAR](https://github.com/dimenwarper/scimitar), the clustering tool [PhenoGraph](https://github.com/jacoblevine/PhenoGraph) (Levine *et al.*, 2015), the single-cell experiment design tool [MIMOSCA](https://github.com/asncd/MIMOSCA) (Dixit *et al.*, 2016), the tree-inference tool [ECLAIR](https://github.com/GGiecold/ECLAIR) (Giecold *et al.*, 2016) and the framework flotilla, which comes with modules for simple visualization, simple clustering and differential expression testing. Hence, only the latter provides a data analysis framework that solves more than one specific task. In contrast to Scanpy, however, flotilla is neither targeted at single-cell nor at large-scale data and does not provide any graph-based methods, which build the core of Scanpy. Also, flotilla is built around a complicated class *Study* that contains data, tools and plotting functions, which orthogonal to the design choice of Scanpy, which is built around a simple class *AnnData* and hence easily extendable (Suppl. Note 3).

## Footnotes

1 DPT compares favorably (Qiu *et al.*, 2017) with Monocle 2 (Trapnell *et al.*, 2014; Qiu *et al.*, 2017), Wanderlust (Bendall *et al.*, 2014) and Wishbone (Setty *et al.*, 2016)

2 The latter uses the JavaScript package [D3.js](http://d3js.org/) for the actual visualization and Python only for preprocessing.

